# Robust taxonomic classification of uncharted microbial sequences and bins with CAT and BAT

**DOI:** 10.1101/530188

**Authors:** F.A. Bastiaan von Meijenfeldt, Ksenia Arkhipova, Diego D. Cambuy, Felipe H. Coutinho, Bas E. Dutilh

## Abstract

Current-day metagenomics increasingly requires taxonomic classification of long DNA sequences and metagenome-assembled genomes (MAGs) of unknown microorganisms. We show that the standard best-hit approach often leads to classifications that are too specific. We present tools to classify high-quality metagenomic contigs (Contig Annotation Tool, CAT) and MAGs (Bin Annotation Tool, BAT) and thoroughly benchmark them with simulated metagenomic sequences that are classified against a reference database where related sequences are increasingly removed, thereby simulating increasingly unknown queries. We find that the query sequences are correctly classified at low taxonomic ranks if closely related organisms are present in the reference database, while classifications are made higher in the taxonomy when closely related organisms are absent, thus avoiding spurious classification specificity. In a real-world challenge, we apply BAT to over 900 MAGs from a recent rumen metagenomics study and classified 97% consistently with prior phylogeny-based classifications, but in a fully automated fashion.

## INTRODUCTION

Metagenomics, the direct sequencing of DNA from microbial communities in natural environments, has revolutionized the field of microbiology by unearthing a vast microbial sequence space in our biosphere, much of which remains unexplored^1–3^. With increases in DNA sequencing throughput, metagenomics has moved from analysis of individual reads to sequence assembly, where increases in sequencing depth have enabled *de novo* assembly of high-quality contiguous sequences (contigs), sometimes many kilobases in length^4^. In addition, current state-of-the-art encompasses binning of these contigs into high-quality draft genomes, or metagenome-assembled genomes (MAGs) ^5–8^. Advancing from short reads to contigs and MAGs allows the field to answer its classical questions^9^, ‘who is there?’ and ‘what are they doing?’ in a unified manner: ‘who is doing what?’, as both function and taxonomy can be confidently linked to the same genomic entity. Because assembly and binning can be done *de novo*, these questions can be applied to organisms that have never been seen before, and the discovery of entirely novel phyla is common still^8^.

Several efficient tools for taxonomic classification of short, read-length sequences have been developed over the years, reflecting the read-based focus of the time. Most tools consider each read as an independent observation, whose taxonomic origin can be estimated by identifying best-hit matches in a reference database, either on read, K-mer, or translated protein level (see ^10^ for an overview). Widely-used programs such as Kraken^11^ (K-mer based), CLARK^12^ (discriminative K-mer based), and Kaiju^13^ (protein-based) can process hundreds of thousands of sequencing reads per second. Without compromising accuracy, still faster approaches use mixture modelling of K-mer profiles, as implemented in FOCUS^14^. Sometimes a Last Common Ancestor (LCA) algorithm is applied to allow for multiple hits with similar scores as the best hit (e.g. Kraken, MEGAN^15^). Kaiju uses a best-hit approach with an LCA algorithm if equally good top-hits are found.

Similar approaches are applied to contigs, with classification often based on the best hit to a reference database. Although fast, the best-hit approach can lead to spurious specificity in classifications, for example when a genomic region is highly conserved or recently acquired via horizontal gene transfer (HGT) from a distantly related organism. As we will show below, the problem is particularly grave when the query contigs are very divergent from the sequences in the database, i.e. they are distantly related to known organisms. Whereas specificity (correctly classified / total classified) can be increased when only classifications at higher taxonomic ranks are considered, this approach is not desirable as taxonomic resolution is unnecessarily lost for query contigs that are closely related to known organisms.

Depending on their length, contigs may contain multiple open reading frames (ORFs), each of which contains a taxonomic signal. Integrating these signals should enable a more robust classification of the entire contig, yet surprisingly few tools exist that integrate distributed signals for contig classification. The viral-specific pipeline MetaVir2^16^ assesses the classification of up to five ORFs encoded on a contig. Recently, the MEGAN long read algorithm was introduced^17^, which partitions the sequence into intervals based on the location of hits of a LAST^18^ search.

In contrast, for taxonomic classification of MAGs it is common to include information from multiple ORFs. Since the classification of complete genomes by using phylogenetic trees of multiple marker genes is well-established^19^, MAG classification has followed these best practices. Some steps in the process can be automated, including initial placement in a low-resolution backbone tree by CheckM^20^, specific marker gene identification and backbone tree taxon selection by phyloSkeleton^21^, and many tools are available for protein alignment, trimming, tree building, and display. However, interpretation of the resulting phylogeny remains a critical manual step, making this approach for genomic taxonomy a laborious task that does not scale well with the increasing number of MAGs being generated (see e.g. ^7^).

Here we present Contig Annotation Tool (CAT) and Bin Annotation Tool (BAT), two tools whose underlying ORF-based algorithm is specifically designed to provide robust taxonomic classification of long sequences and MAGs that contain multiple protein-encoding gene sequences. Both tools require minimal user input and can be applied in an automated manner, yet all aspects are flexible and can be tuned to user preferences.

## Benchmarking classification of sequences from novel taxa

Taxonomic classifiers are often benchmarked by testing them on sequences from novel taxa, i.e. that are not (yet) in the reference database (e.g. ^11,12,14^). Alternatively, unknown query sequences can be simulated by using a ‘leave-one-out’ approach, where the genome that is being queried is removed from the database (e.g. ^13,17^). However, due to taxonomic biases in the database composition, other strains from the same species, or other species from the same genus may still be present. Thus, the leave-one-out approach does not reflect the level of sequence unknownness that is often encountered in real metagenomes, where the query sequences may be only distantly related to the ones in the reference database. Here, we rigorously assess the performance of contig classification tools by developing an extensive database reduction approach at different taxonomic ranks, where novel species, genera, and families are simulated by removing all the sequences of entire taxa from the database. We show that the algorithm of CAT and BAT allows for the correct classification of organisms from known and unknown taxa, at least up to the rank of novel families. We compared the performance of CAT to contig classifications by a best-hit approach, LAST+MEGAN-LR^17^, and Kaiju^13^. Moreover, we used BAT to classify a large, recently published set of 913 MAGs from the cow rumen^7^, and whose taxonomic classifications involved extensive phylogenetic analyses. We show that CAT and BAT perform in par with, or better than existing methods, especially for sequences that are highly unknown.

## RESULTS AND DISCUSSION

### Contig classification with CAT

#### CAT has two user definable parameters

We used CAT (Figure 1) to classify ten simulated contig sets in the context of four reference databases with different levels of simulated unknownness, representing query sequences from (A) known strains, (B) novel species, (C) novel genera, and (D) novel families (see Online methods). To assess the effect of the two key user parameters, *r* (hits included within *range* of top hits) and *f* (minimum *fraction* classification support) on precision, fraction of classified contigs, sensitivity, and taxonomic rank of classification, we ran CAT with a wide range of possible parameter values against all four reference databases (Figure 2). This parameter sweep revealed a trade-off between the classification precision on the one hand, and the taxonomic resolution and the fraction of classified contigs on the other. This general trend can be understood by considering that classifications at a low taxonomic rank (i.e. close to the species level, high taxonomic resolution) will inevitably be increasingly imprecise, especially if closely related organisms are absent from the reference database. This might be resolved by classifying contigs at a higher taxonomic rank, but this leads to increased numbers of contigs not being classified or classified at trivially informative taxonomic ranks such as ‘cellular organisms’ or ‘root’.

**Figure 1.**
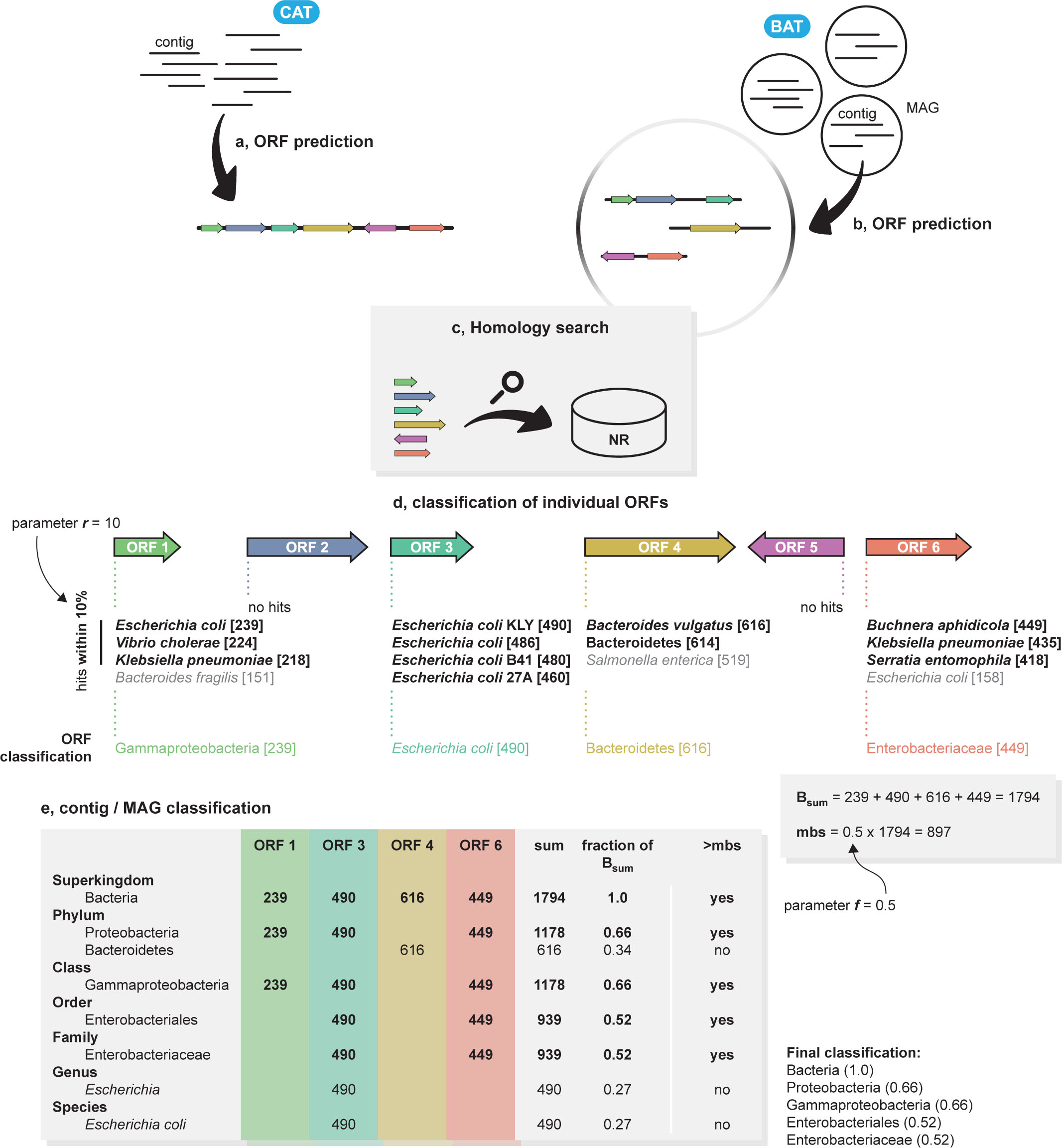
Contig and MAG classification with CAT and BAT. (**a, b**) Step 1: ORF prediction with Prodigal. CAT analyses all ORFs on a contig, BAT analyses all ORFs in a MAG. (**c**) Step 2: predicted ORFs are queries with Diamond to the NCBI non-redundant protein database (NR). (**d**) Step 3: ORFs are individually classified based on the LCA of all hits falling within a certain range of the top hit (parameter *r*), and the top-hit bit-score is assigned to the classification. Bit-scores of hits are depicted within brackets. Hits in grey are not included in final annotation of the ORF. Parameter *f* defines minimum bit-score support (mbs). (**e**) Step 4: contig or MAG classification is based on a voting approach of all classified ORFs, by summing all bit-scores from ORFs supporting a certain classification. The contig or MAG is classified as the lowest classification reaching mbs. The example illustrates the benefit of including multiple ORFs when classifying contigs or MAGs; a best-hit approach might have selected *Bacteroides vulgatus* or Bacteroidetes if an LCA algorithm was applied as its classification, as this part has the highest score to proteins in the database in a local alignment-based homology search. In the example only six taxonomic ranks are shown for brevity, in reality CAT and BAT will interpret the entire taxonomic lineage.

**Figure 2.**
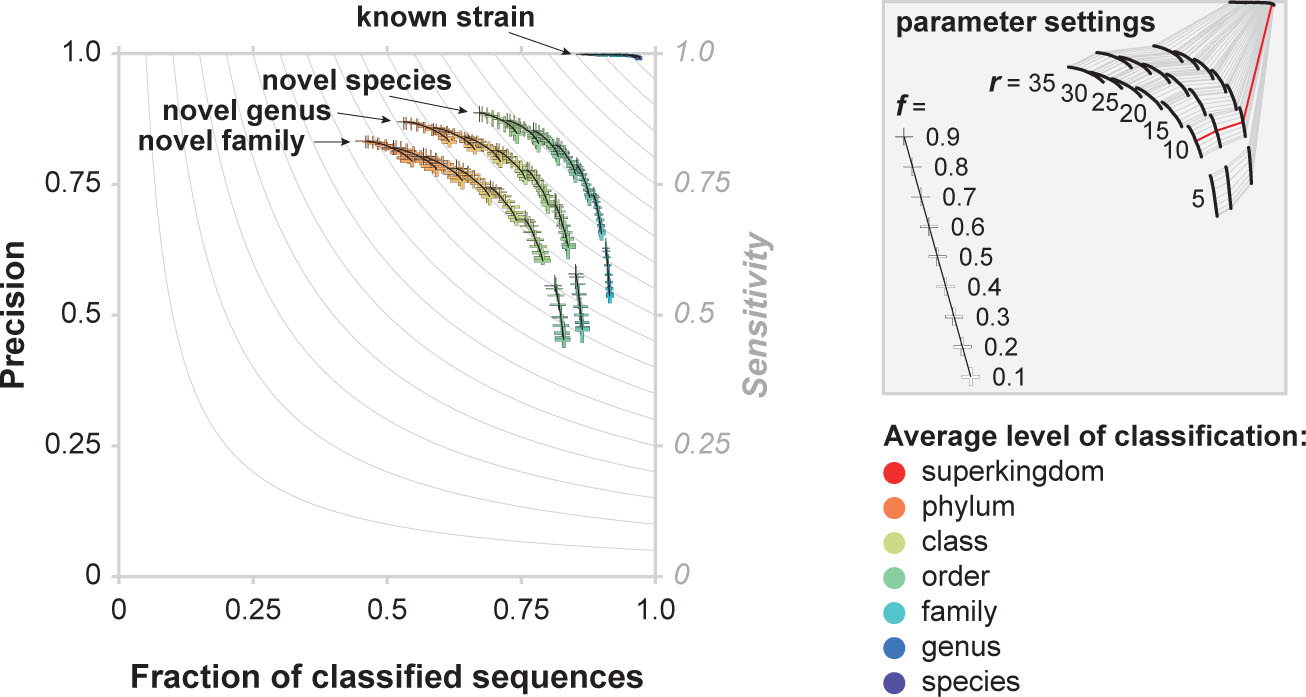
Classification performance of CAT for different levels of unknownness across a range of parameter settings. Thickness of markers indicate values of the *f* parameter, runs with similar *r* parameter values are connected with black lines. Markers indicate maximum and minimum values out of ten benchmarking datasets, bars cross at the average. Colour coding indicates average taxonomic rank of classification across the then benchmarking dataset (minimum and maximum values not shown). Grey lines in the plot depict sensitivity, which is defined as fraction of classified contigs x precision. Runs with equal parameter settings are connected in the parameter settings figure, showing that CAT achieves a high precision regardless of unknownness of the query sequence, by classifying sequences that are more unknown at higher taxonomic ranks. Default parameter combination (*r* = 10, *f* = 0.5) is shown in red.

The *r* parameter, which governs the divergence of included hits for each ORF, had the largest effect. As increasing *r* includes homologs from increasingly divergent taxonomic groups, their LCA is pushed back and classifications at low taxonomic ranks are lost, resulting in fewer classified contigs at lower taxonomic resolution (higher ranks), but with higher average precision. The *f* parameter, which governs the minimum bit-score support required for classifying a contig, has a smaller effect. Decreasing *f* allows CAT classifications to be based on evidence from fewer ORFs, leading to more tentative classifications at lower taxonomic ranks. As a result, more contigs are classified at lower taxonomic ranks, albeit with a lower precision.

As a user increases *r* and *f*, this will increasingly result in high rank classifications that are correct but ultimately uninformative. When low values of *r* and *f* are chosen, the classifications will be more specific (i.e. at a lower taxonomic rank) but more speculative (i.e. precision goes down). Based on the parameter sweep described above, we set the default values for CAT contig classification to *r* = 10; *f* = 0.5 (red line in the legend of Figure 2). Note that this value of *f* = 0.5 results in at most one classification at a given taxonomic rank, since >50% of the bit-score supports that classification.

#### Comparison to state-of-the-art tools

We compared classification by CAT to (1) the recently published LAST+MEGAN-LR algorithm^17^, (2) the widely used Kaiju algorithm^13^, and (3) a standard best-hit approach with Diamond^22^ (Figure 3). Kaiju was designed for short-read classification, but its underlying algorithm allows for the classification of long sequences as well, and has recently been used as such^17,23,24^. Final classification is based on the hit with the maximum exact match (MEM), or on the highest scoring match allowing for mismatches (Greedy).

**Figure 3.**
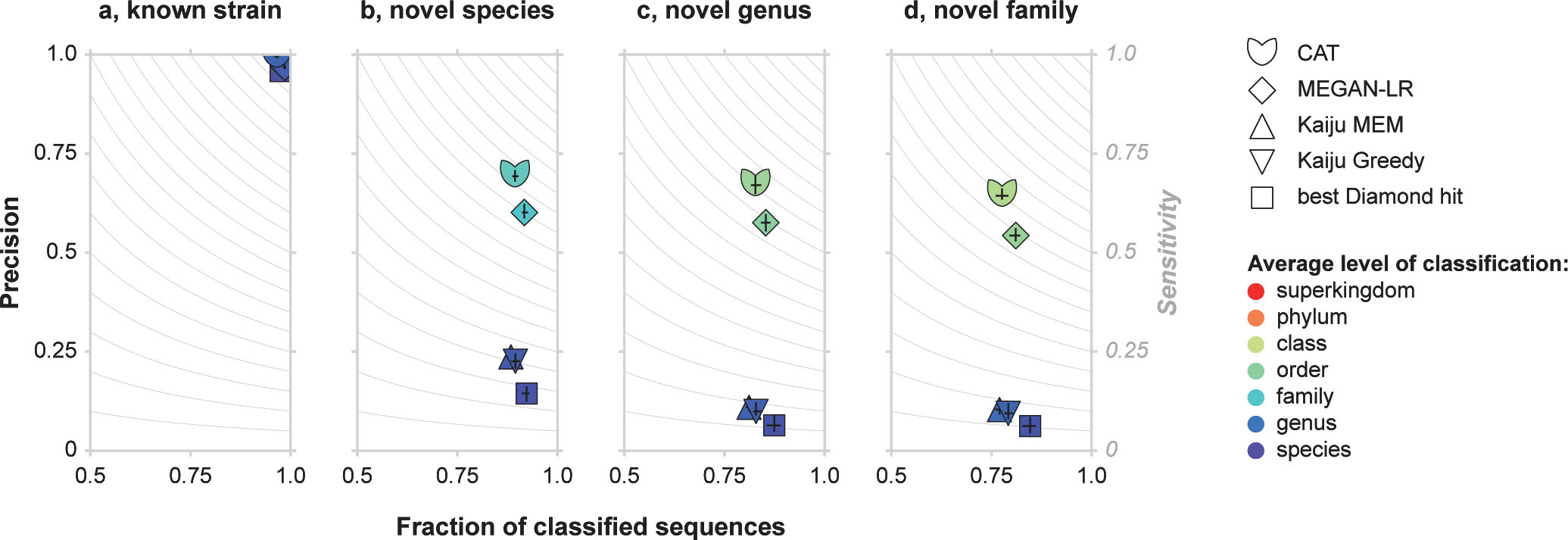
Classification performance of CAT, LAST+MEGAN-LR, Kaiju, and Diamond best-hit for different levels of unknownness. (**a**) Classification of known sequences, (**b-d**) classification of simulated novel taxa for different levels of divergence from reference databases. Black bars indicate maximum and minimum values out of ten benchmarking datasets, bars cross at the average. Colour coding indicates average taxonomic rank of classification across the then benchmarking dataset (minimum and maximum values not shown).

When classifying simulated contigs against the full reference database (known strains), all programs showed a similar precision and fraction of classified sequences (Figure 3A). The average taxonomic rank of classification is slightly higher for CAT (0.77 ± 0.05) and LAST+MEGAN-LR (0.62 ± 0.04) than for best-hit (0.14 ± 0.03), Kaiju MEM (0.25 ± 0.04) and Kaiju Greedy (0.25 ± 0.04), reflecting the conservative LCA-based classification strategies of the former two. Best-hit does not use LCA, and Kaiju only in cases were multiple hits have identical scores, and thus they classify contigs according to the taxonomic rank of their match in the reference database.

When novel species, genera, and families were simulated by removing related sequences from the database, precision declined rapidly for best-hit and Kaiju (Figure 3A-D). The classifications called by these approaches are often too specific, because in databases where closely related sequences are absent, the singular best hit may still match a sequence that is annotated at a low taxonomic rank, although this annotation cannot match that of the query. This spurious specificity can be seen in the average rank of classification, which stays close to the species rank, even when sequences from the same species, genus, or family were removed from the database (Figure 3B-D). CAT and LAST+MEGAN-LR clearly perform better in the face of such uncharted sequences. With default parameter settings, CAT has higher precision and sensitivity than MEGAN-LR and classifications are made at slightly higher taxonomic ranks, e.g. for simulated novel families (Figure 3D) 3.40 ± 0.07 and 3.01 ± 0.07 for CAT and MEGAN-LR, respectively.

#### CAT automatically classifies sequences at the appropriate taxonomic rank

As a solution to the spurious specificity of the best-hit approach described above, best-hit classifications may be assigned to a higher taxonomic rank, such as genus, family, or even phylum. Supplementary Figure 2 shows that application of a rank cut-off to the best-hit classifications only partly solved the problem of spurious specificity. For instance, when only order rank classifications were considered, precision was 0.21 ± 0.02 in the case of simulated novel families (Supplementary Figure 2D). Importantly, applying a rank cut-off may unnecessarily sacrifice taxonomic resolution in cases where the query sequences do have close relatives in the reference database and classification at a low taxonomic rank would be justified.

As shown in Figure 2, the ORF-based voting algorithm of CAT ensures a high precision regardless of the level of unknownness of the query sequences, i.e. whether closely related sequences are present in the reference database or not. Only when necessary, taxonomic resolution is traded for precision: when classifying contigs that are more distantly related to the sequences in the reference database, hits will have weaker bit-scores and match sequences that are taxonomically more diverse. As a result of these conflicting signals, the CAT algorithm automatically increases the taxonomic rank when classifying more divergent query sequences. For example, when novel families are simulated, precision is 0.64 ± 0.008 and the average taxonomic rank of classifications lay between order and class (Supplementary Figure 2D). In contrast, best-hit classifications that were cut-off at these taxonomic ranks had a precision of 0.21 ± 0.02 and 0.47 ± 0.01, respectively, with a similar fraction of classified sequences.

#### CAT is fast and has a very low memory requirement

CAT is about two times faster than LAST+MEGAN-LR (Figure 4A) and outperforms all other programs in terms of memory-usage. The most memory intensive step is the Diamond search for homologs in the vast NR database (Figure 4B). Note that construction of the database files during the ‘prepare’ step requires more memory than the classification, but this only needs to be done once.

**Figure 4.**
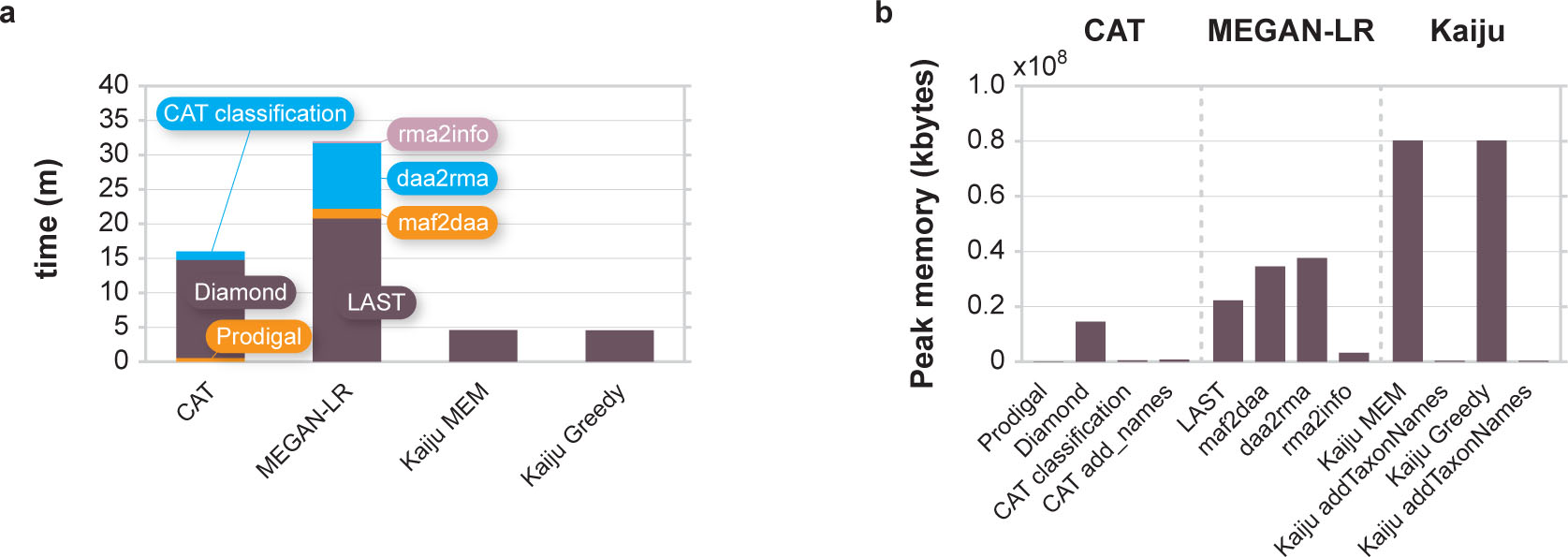
Computer resource usage by CAT, LAST+MEGAN-LR, and Kaiju. (**a**) Run-time and (**b**) peak memory usage. In a, classification by CAT and Kaiju include adding taxonomic names to the classification, in b these steps are depicted separately.

### Bin classification with BAT

#### Classification of 913 Metagenome-Assembled Genome bins (MAGs)

Next, we set out to apply the algorithm to MAGs, i.e. draft genomes that can be generated from metagenomes by assembly and binning. Since the typical pipeline to generate MAGs is reference database independent, they can be distantly related to known organisms. As benchmark set, we picked 913 recently published MAGs from the cow rumen^7^ that represented a wide range of novelty at different taxonomic ranks (Supplementary Figure 3A). The published classifications were based on placement of the MAGs in a backbone tree and subsequent refinement, a slow process that includes various manual steps and visual screening^7^. At the time of our study, the MAGs were not yet included in the reference database, providing an ideal test case for our automated classification tool BAT.

The 913 MAGs were previously assessed to be ≥80% complete, have ≤10% contamination, and contain between 541-5,378 ORFs each (Supplementary Figure 3B). We ran BAT with default parameter settings for MAGs classification (*r* = 5, *f* = 0.3). The low *r* value ensures that individual ORFs are annotated to a LCA with a relatively low taxonomic rank, as hits within 5% of the highest bit-score are considered. The low *f* value reports taxonomic classifications that are supported by at least 30% of the bit-score evidence. While this could be considered a speculative call when contigs with relatively few encoded ORFs are annotated, the much higher number of ORFs in MAGs means that even classifications with relatively low *f* values are backed by a high number of ORFs. We scored the consistency between BAT and the published classifications (Figure 5A), dividing consistent classifications into three groups: (i) BAT can be more conservative than the published classification, i.e. BAT classifies the MAG to an ancestor of the published classification, (ii) classifications can be equal, and (iii) BAT can be more specific. Alternatively, BAT can classify a MAG inconsistently, i.e. in a different taxonomic lineage than the original publication. As shown in Figure 5A, 885 of 913 MAGs (97%) were classified consistently with the original publication. If parameter *f* is relaxed, average level of classification for the MAGs increases (Figure 5B). Importantly, decreasing the value of *f* has little effect on inconsistency rate. Thus, changing this parameter will mainly lead to a change in the rank of classification, while the taxonomic lineage will remain unchanged.

**Figure 5.**
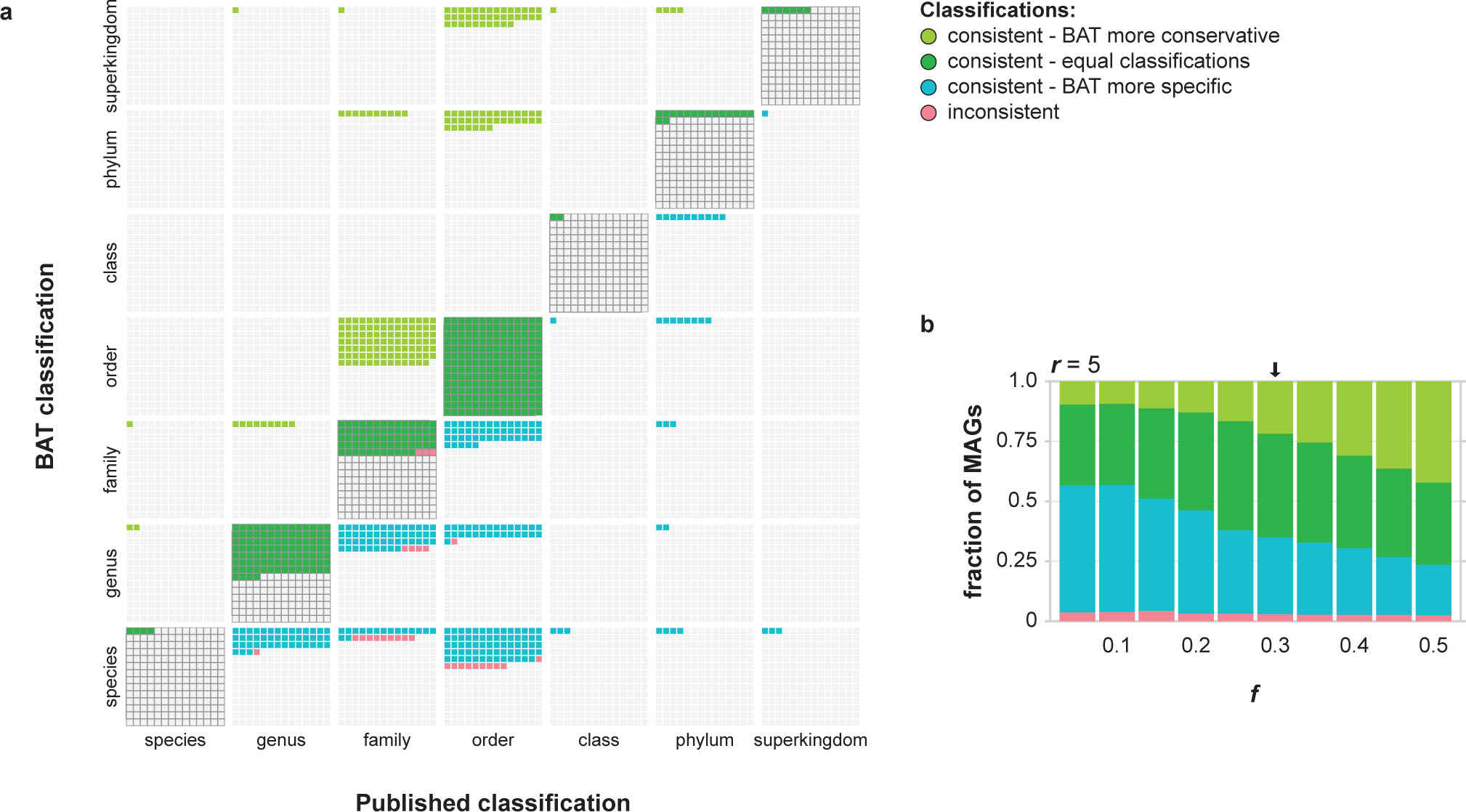
Classification of 913 MAGs with BAT. (**a**) Consistency between BAT classifications and published classifications with default parameter settings (*r* = 5, *f* = 0.3). (**b**) The average level of classification can be increased by increasing *f*. Arrow indicates BAT results for its default parameter settings.

To assess the taxonomy of the 28 inconsistently classified MAGs (at r = 5, f = 0.3), we placed them in a phylogenomic tree with closely related genomes and observed their closest relatives, the published classifications, and the BAT classifications. As shown in Figure 6, BAT classified all 28 inconsistent MAGs more precisely and at a higher taxonomic resolution than the published classifications. Note that this may be due to these closely related reference genomes being new additions to the database since the research was performed. Together, these results highlight the benefit of using BAT for the rapid, automated, and high resolution taxonomic classification of novel microbial lineages at a range of unknownness.

**Figure 6.**
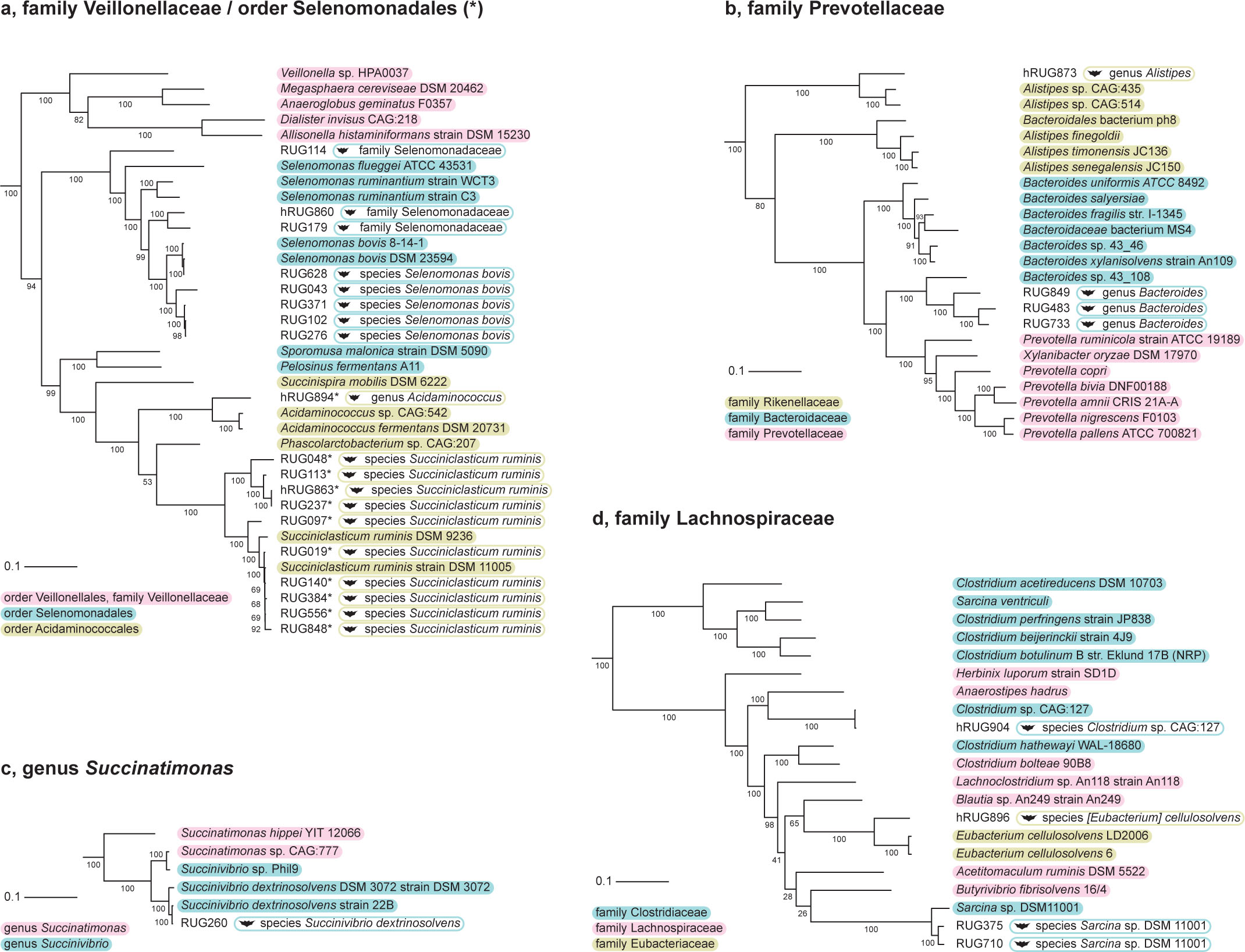
Tree placement of the 28 inconsistently classified MAGs that were assigned to five different taxa according to the original classifications (**a-d**). Headers of subfigures refer to the published classifications. In a, MAGs published as Selenomonadales are marked with an asterisk. Taxonomic classification of reference genomes is indicated in shades boxes. BAT classifications of MAGs are indicated in open boxes.

## CONCLUSIONS

Metagenomics continues to reveal novel microorganisms in all environments in the biosphere, whose sequences can be reconstructed with high accuracy by using high-throughput DNA sequencing and modern sequence assembly tools. Taxonomically classifying these uncharted sequences remains challenging, partly because the vast natural biodiversity remains highly under-represented in even the largest reference databases, and partly because existing tools are built to classify short sequencing reads.

We presented CAT and BAT, a suite of tools that exploit database searches of individual ORFs, LCA annotation, and a voting algorithm to classify long contigs and metagenome-assembled genomes (MAGs). As we have shown, these query sequences contain a wealth of information that allows their accurate taxonomic classification at appropriate taxonomic ranks, i.e. at a low rank when closely related organisms are present in the database, and at a high rank when the sequences are divergent or highly novel. We have shown that the low precision of conventional best-hit approaches when classifying novel taxa can be overcome by a voting algorithm based on classifications of multiple ORFs. Elegantly, sequences from organisms that are distantly related to those in the reference database are automatically classified at a higher taxonomic rank than known strains. ORFs on divergent sequences will hit a wider variety of different taxa both on the individual ORF level and between ORFs. Such conflict of classifications is automatically resolved by the algorithm by providing a more conservative classification, so no taxonomic cut-off rank for classification needs to be pre-defined. In metagenomes containing both known and unknown sequences, the algorithm vastly outperforms best-hit approaches in both precision and sensitivity.

CAT and BAT supplement a modern metagenomics workflow in various ways. For example, CAT can be used after metagenome assembly to confidently classify all contigs. We expect that subsequent classification of the original sequencing reads in terms of these contigs will result in fewer false positives than if reads were taxonomically classified using short-read classification tools. Moreover, BAT will rapidly provide taxonomic classifications of MAGs without requiring a full phylogenomics pipeline and subsequently visual inspection of the tree. CAT classifications of individual contigs within MAGs can be used to identify taxonomic outliers, and flag those as possible contamination. As most binning tools do not incorporate taxonomic signals (e.g. ^25,26^), CAT classification can be considered as independent evidence and might be used to decide on the inclusion of specific contigs in a MAG.

BAT provides a robust and rapid classification of MAGs in a single operation, but it is not a replacement for high-confidence phylogenomic tree construction based on marker gene superalignments which remains the gold standard^19^. However, BAT queries the full NCBI non-redundant reference database (NR) and the taxonomic context is thus much bigger than any phylogenomic tree that depends on completely sequenced genomes. For example, the backbone tree of CheckM currently includes only 5,656 genomes^20^. BAT classification is fully automated and can be run on a set of MAGs with minimal user input, allowing MAG classification to be scaled up considerably as we showed here for over nine hundred MAGs that were classified consistently with the original publication in almost all cases.

As long as sequence space is incompletely explored and reference databases represent a biased view of the tree of life^1,3^, algorithms designed to address the abundant uncharted microbial sequences will be needed to make sense of the microbial world. Decreasing sequencing costs and improvement of alignment and binning algorithms have moved metagenomics from the analysis of short reads towards contigs and MAGs, improving our understanding of microbial ecosystems to a genomic resolution. As these data will only increase in the coming years, we presented a robust solution to their specific challenges that we expect will play an important role in future metagenomics workflows.

## AVAILABILITY

CAT and BAT are available at https://github.com/Dutilh/CAT/. All benchmarking datasets and reference databases with increasing levels of unknownness are available from the authors upon request.

## SUPPLEMENTARY DATA

Methods and Supplementary Figures are available online.

## AUTHOR CONTRIBUTIONS

B.E.D. conceived the study. D.D.C., K.A., and F.A.B.v.M. wrote the code. F.H.C. and F.A.B.v.M. analysed the data. F.A.B.v.M. and B.E.D. wrote the paper. All authors read and approved the manuscript.

## Online Methods

### Software availability

CAT and BAT are implemented in Python 3. The software and user manual are available at https://github.com/Dutilh/CAT/.

### Explanation of the algorithm

Both CAT and BAT take high-quality long DNA sequences in FASTA format as input (Figure 1), such as assembled contigs or corrected long Nanopore or PacBio reads^1, 2^. First, ORFs are predicted with Prodigal^3^ in metagenome mode, using default parameter settings (genetic code 11) (Figure 1A, B). Predicted proteins can also be independently supplied to CAT/BAT in case a user prefers a different gene caller than Prodigal.

Next, protein translations of the predicted ORFs are queried against the National Center for Biotechnology Information (NCBI) non-redundant protein database (NR) ^4^ using Diamond^5^ blastp (e-value cut-off of 0.001, BLOSUM62 alignment matrix, reporting alignments within 50% range of top hit bit-score) (Figure 1C). The NR database is currently the largest sequence database where all sequences are assigned to clades in NCBI Taxonomy^6^. A separate Blast tabular output file can also be supplied together with the predicted proteins file, in which case CAT/BAT starts directly with classification.

Taxonomic classification of the query sequences is then carried out based on a voting approach that considers all ORFs on a query with hits to the reference database. Here, the main difference between CAT and BAT is that CAT considers ORFs on a single contig, whereas BAT considers ORFs on all contigs belonging to a MAG. CAT and BAT also have slightly different default parameters values (see below).

First, the algorithm infers the taxonomic affiliation of individual ORFs based on the top Diamond hits (Figure 1D). To account for similarly high-scoring hits in potentially different clades, hits within a user-defined range of the top hit bit-score to that ORF are considered and the ORF is assigned to the LCA of their lineages (parameter *r* for *range*, by default hits with bit-scores within 10% or 5% range of the top hit bit-score are included, *r* = 10 for CAT and *r* = 5 for BAT, respectively). By adjusting parameter *r*, the user can tune how conservative CAT is in the classification of individual ORFs. For example, increasing *r* results in more divergent hits being included that together are likely to have a deeper LCA, thus leading to a more conservative ORF classification at a higher taxonomic rank. In contrast, decreasing *r* leads to a more specific classification since fewer and more similar hits will be included, likely with a narrower taxonomic range. This accounts for conserved or HGT-prone genes that are highly similar in diverse taxa by assigning them a high-rank classification. The top hit bit-score for each ORF is registered for the subsequent voting process (Figure 1D).

Next, the query contig or MAG is evaluated by summing the bit-scores for each taxon identified among the classifications of all ORFs, as well as their ancestral lineages up to the taxonomy root (Figure 1E). The query contig or MAG is then assigned to a taxon, if the total bit-score evidence for that taxon exceeds a cut-off value (*mbs*, minimum bit-score support), which is calculated as a fraction (parameter *f* for *fraction*) of the sum of the bit-scores of all ORFs (mbs = *f* * B_sum_, by default *f* = 0.5 for CAT and *f* = 0.3 for BAT). For example, if parameter *f* is set to 0.5, this means that a contig is assigned to a taxon if at least half of the sum of the bit-scores of all ORFs supports that classification (mbs = 0.5 × B_sum_). This is done at all taxonomic ranks including phylum, class, order, family, genus, and species. The algorithm stops at the taxonomic rank where the total bit-score supporting the classification drops below the minimum bit-score support value, so CAT/BAT automatically finds the lowest rank taxonomic classification that is still reliable (Figure 1E). Note, that if *f* < 0.5, multiple lineages at a given taxonomic rank may exceed the threshold, and all are written to the output file.

### Output files

For each query contig or MAG, the full taxonomic lineage of the lowest-rank supported classification is written to the output file, together with support values (i.e. the fraction of B_sum_ that is represented by the taxon). In addition, the number of ORFs found on the contig or MAG and the number of ORFs on which the classification is based are written to the output file. An extra output file containing information about individual ORFs is also generated, including classifications of ORFs and an explanation for any ORF that is not classified. We advise the user caution when interpreting the classifications of short contigs that are based on relatively few ORFs as they will be less robust than the classifications of long contigs or MAGs.

### Helper programs

The CAT/BAT package comes bundled with three helper utilities, ‘prepare’, ‘add_names’, and ‘summarise’. ‘Prepare’ only needs to be run once. It downloads all the needed files including NCBI taxonomy files and the NR database. It constructs a Diamond database from NR, and generates the files needed for subsequent CAT and BAT runs. Because the first protein accession in NR not always represents the LCA of all protein accessions in the entry, ‘prepare’ corrects for this in the protein accession to taxonomy id mapping file (prot.accession2taxid). After running CAT/BAT, ‘add_names’ will add taxonomic names to the output files, either of the full lineage or of official taxonomic ranks alone. ‘Summarise’ generates summary statistics based on a named classification file. For contig classification, it reports the total length of the contigs that are classified to each taxon. For MAG classification, it reports the number of MAGs per taxon.

### Generation of contig benchmarking datasets

To test the performance of CAT, we artificially generated contigs from known genome sequences in the RefSeq database^7^. We randomly downloaded one genome per taxonomic order from bacterial RefSeq on July 7^th^, 2017 (163 orders in total) and cut the genomes into at most 65 non-overlapping contigs, generating a set of ~10,500 contigs with known taxonomic affiliation. Contig lengths were based on the length distribution of eight assembled real metagenomes deposited in the Sequence Read Archive (SRA)^8^ (assembly with metaSPAdes v3.10.1^9^ after quality filtering with BBDuk that is included with BBTools v36.64 (https://sourceforge.net/projects/bbmap/), see Supplementary Table 1), with a minimum length of 300 nucleotides. This was done ten times to construct ten different benchmarking datasets sampled from 163 different genomes, each from a different taxonomic order (Supplementary Table 1).

Viruses remain vastly under-sampled and the sequences in the database remain a small fraction of the total viral sequence space^10^. Moreover, the hierarchy of the viral taxonomy is not as deeply structured as the taxonomy of cellular organisms^11^. Based on these considerations, we did not explicitly assess the performance of our tool on viral sequences. However, we expect that classification of viruses will be readily possible when closely related viruses are present in the reference database.

### Reference databases with increasing levels of unknownness

The benchmarking datasets generated above are derived from genomes whose sequences are also present in the reference database, corresponding to the perhaps unlikely scenario where the query sequences in the metagenome are identical to known strains in the database. To benchmark our tools in the context of discovering sequences from novel taxa, we next generated novel reference databases with increasing levels of unknownness by removing specific taxonomic groups from NR. In addition to the original NR database (known strains), three derived databases were constructed to reflect the situation of discovering novel species, genera, and families. This was done by removing all proteins that are only present in the same species, genus, or family as any of the 163 genomes in the benchmarking dataset. To do this, we either removed the sequences from the database itself, or if a protein was identical in sequence to a protein in another clade, we changed the protein accession to taxonomy id mapping file to exclude the query taxon. In contrast to many other taxonomic classification tools, all the programs that we compared (CAT, Diamond best-hit, LAST+MEGAN-LR, and Kaiju) allowed such custom files to be used. The three reduced databases and associated mapping files thus reflect what NR would have looked like if the species, genus, or family of the genomes present in the benchmarking dataset were never seen before. This was done independently for each of the ten different benchmarking datasets, resulting in a total of 30 new reference databases to rigorously test the performance of our sequence classification tools in the face of uncharted microbial sequences. Simulating unknownness like this provides a better benchmark for classification of unknown sequences than a leave-one-out approach where only the query genome is removed from the reference database^12, 13^, because close relatives of the query may still be present. It also avoids the need to simulate the sequences themselves, and the many assumptions associated with it^14^.

### Programs, parameters, and dependencies

NR database and taxonomy files were downloaded on the November 23^rd^, 2017. Prodigal v2.6.3^3^ was used to identify ORFs on the simulated contigs. Diamond v0.9.14^5^ was used to align the encoded proteins to the reference databases for CAT and for the Diamond best-hit approach. Kaiju v1.6.2^12^ was run both in MEM and Greedy mode with SEG low complexity filter enabled. The number of mismatches allowed in Greedy mode was set to 5. For LAST+MEGAN-LR, LAST v914^15^ was used to map sequences to the databases with a score penalty of 15 for frame-shifts, as suggested in^13^. Scripts in the MEGAN v6.11.7^13^ tools directory were used to convert LAST output to a classification file. The maf2daa tool was used to convert LAST output to a .daa alignment file. The daa2rma tool was used to apply the long read algorithm. ‘--minSupportPercent’ was set to 0, the LCA algorithm to longReads, and the longReads filter was applied. ‘--topPercent’ was set to 10 and ‘--lcaCoveragePercent’ to 80 (MEGAN-LR defaults). The rma2info tool was used to convert the generated .rma file to a classification file. When a reduced database was queried, the appropriate protein accession to taxonomy id mapping file was supplied via its respective setting (see section ‘Reference databases with increasing levels of unknownness’ above).

### Scoring of contig classification performance

For contig classification, we scored (i) the fraction of classified contigs, (ii) sensitivity, (iii) precision, and (iv) average rank of classification (Supplementary Figure 1). Classifications were compared at the taxonomic ranks of species, genus, family, order, class, phylum, and superkingdom. In those cases where *f* < 0.5 and multiple classifications reached the mbs threshold, we chose the lowest classification that reached a majority vote (i.e. as if *f* = 0.5) for calculating the four performance measures i-iv. This means CAT classifications were more conservative in those (rare) cases. Contigs with a classification higher than the superkingdom rank (e.g. ‘cellular organisms’ or ‘root’) were considered unclassified, as these classifications are trivially informative in our benchmark. For all tools, a classification was considered correct if it was in the true taxonomic lineage, regardless of rank of classification. The average taxonomic rank of classification was calculated for all classified contigs, where the ranks species-phylum were given the integer values 0-6, respectively. Sensitivity and precision were scored as (correctly classified / total number of contigs) and (correctly classified / total number of classified contigs), respectively.

### Bin classification

913 high-quality draft genome bins (MAGs) (completeness ≥80%, contamination ≤10%) from the cow rumen generated with both conventional metagenomics as well as Hi-C binning methods^16^ were downloaded from the DataShare of the University of Edinburgh (https://datashare.is.ed.ac.uk/handle/10283/3009). Taxonomic classification of the MAGs was downloaded from the Supplementary Data that accompanies the paper, and manually corrected if the names did not match our taxonomy files. To save disk space on the alignment file being generated, we ran BAT on batches of 25 genomes each. Akin to the contig classification case, we only considered classifications by BAT at official taxonomic ranks, and chose the majority classification in those cases were BAT gave more than one classification for a MAG (i.e. as if *f =* 0.5 for that MAG) resulting in more conservative classifications.

To manually assess the 28 MAGs that whose classification was inconsistent with the published classifications, we created a phylogenomic tree of those bins together with closely related genomes that were downloaded from PATRIC^17^ on January 16^th^, 2018. CheckM v1.0.7^18^ was used to extract 43 phylogenetically informative marker genes that were realigned with ClustalOmega v1.2.3^19^. We concatenated the alignments to create a superalignment and included gaps if a protein was absent. We constructed a maximum likelihood tree with IQ-TREE v1.6.3^20^, with ModelFinder^21^ set to fit nuclear models (best-fit model LG+R7 based on Bayesian Information Criterion), including 1,000 ultrafast bootstraps^22^. Per clade, rooted subtrees were visualized in iTOL^23^.

### Usage of computer resources

Run time and peak memory usage were estimated with the Linux /usr/bin/time utility. Elapsed wall clock time and maximum resident set size were scored for a run classifying contig set #1 (10,533 contigs, see Supplementary Table 1) with the NR reference database. All tools were run with default parameter settings. Runs were performed on a machine with an Intel Xeon Gold 6136 Processor, 125.6 GB of memory, 24 cores and 48 threads. Whenever one of the programs allowed for the deployment of multiple threads, all were used. CAT and BAT have been tried and tested on 125 GB machines.

## Table and Figures Legends

**Supplementary Figure 1.**
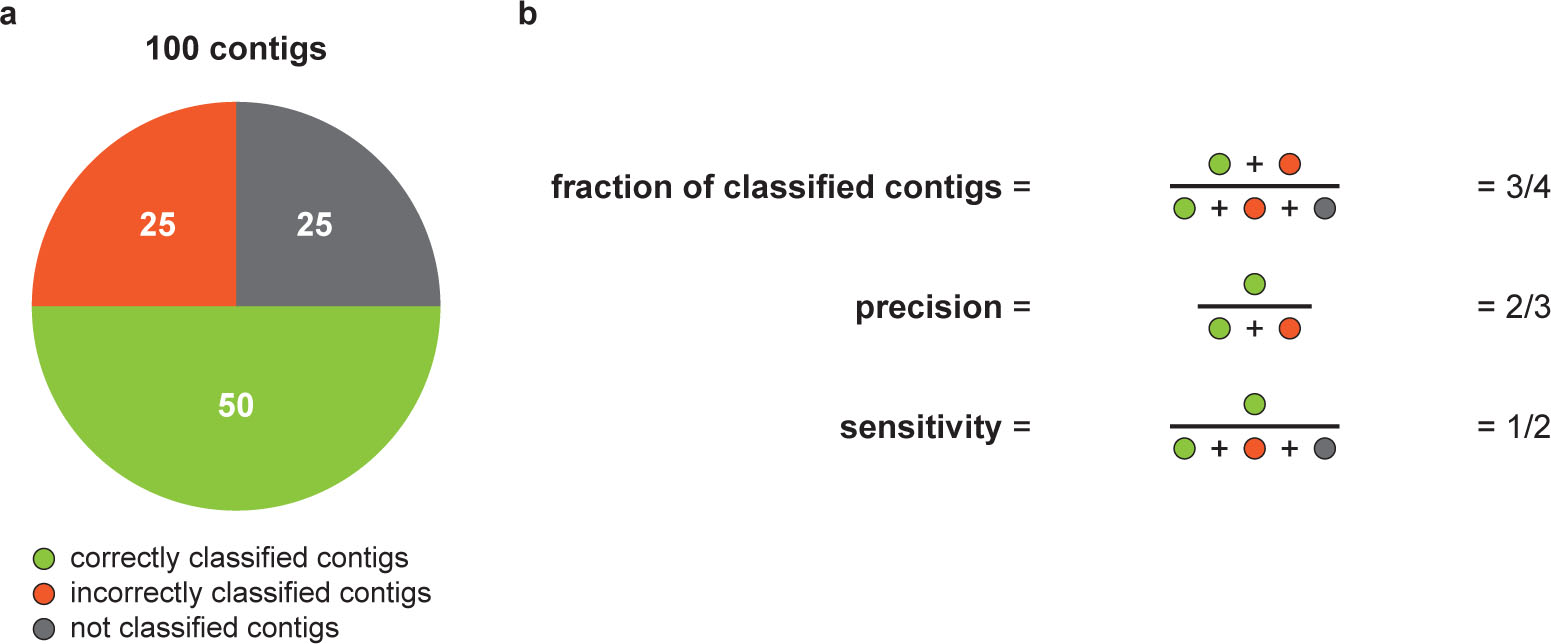
Measuring performance for contig classification. (**a**) Example contig set. Classifications above superkingdom rank (e.g. ‘cellular organisms’ or ‘root’) are considered not classified. Half of the total classifications is contained within the true taxonomic lineage and is thus scored as correct, and a quarter is not. (**b**) Measures of performance. Sensitivity is fraction of classified contigs x precision. Average taxonomic rank of classification is calculated for all classified contigs (75 in the example), where the ranks species-phylum are given the integer values 0-6, respectively, allowing an average to be calculated.

**Supplementary Figure 2.**
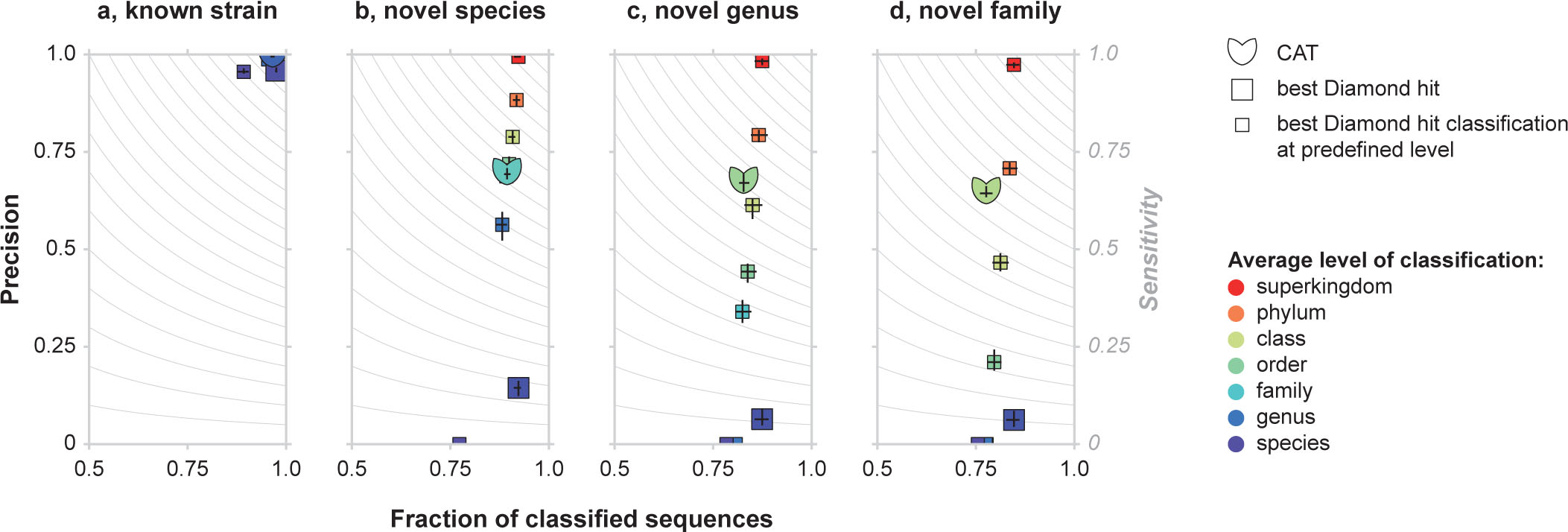
Classification performance of CAT, Diamond best-hit, and Diamond best-hit with different taxonomic rank cut-offs. (**a**) Classification of known sequences, (**b-d**) classification of simulated novel taxa for different levels of divergence from reference databases. Black bars indicate maximum and minimum values out of ten benchmarking datasets, bars cross at the average. Colour coding indicates average taxonomic rank of classification across the then benchmarking dataset (minimum and maximum values not shown).

**Supplementary Figure 3.**
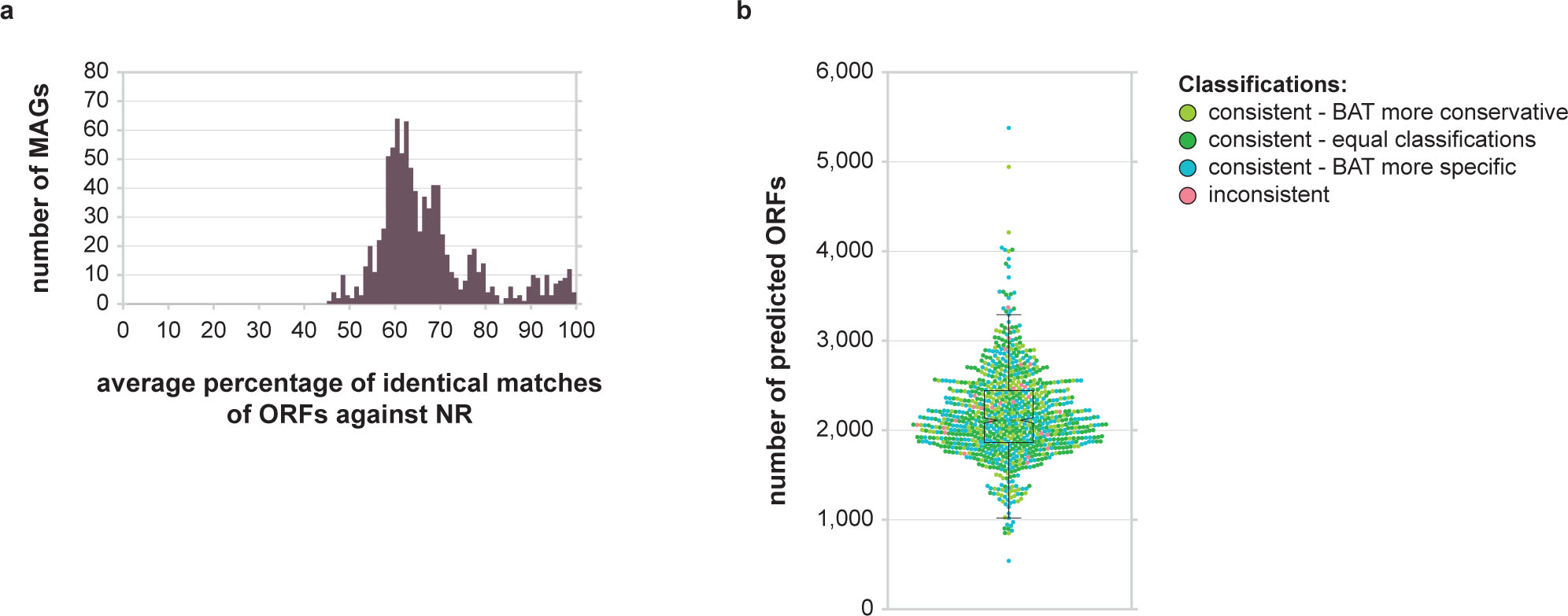
Predicted ORFs on 913 MAGs. (**a**) Average percentage of identical matches with the best Diamond hit in the NR database for all predicted ORFs in a MAG. The wide distribution shows that the MAGs represent a wide range of novelty, i.e. most MAGs are organisms that are not present in the NR database yet. (**b**) Swarmplot showing the number of predicted ORFs per MAG. MAGs are coloured as in Figure 6 (*r* = 5, *f* = 0.3).

Supplementary Table 1. Ten benchmarking contig sets were generated from genomes deposited in bacterial RefSeq. Lengths were based on the length distribution of eight assembled real metagenomes deposited in SRA (libraries SRR2922420, ERR1198954, ERR315808, ERR315819, SRR3666246, ERR594326, ERR599045, SRR3732372). Reads were quality filtered with BBDuk (BBTools v36.64), and assembled with metaSPAdes v3.10.1. Contigs had a minimum length of 300 nucleotides. RefSeq id, length, start and stop coordinate in the genome, and taxonomic classifications of contigs are shown. Datasets are available from the authors upon request.

